# Identification of SGLT2 inhibitor Ertugliflozin as a treatment for COVID-19 using computational and experimental paradigm

**DOI:** 10.1101/2021.06.18.448921

**Authors:** Shalini Saxena, Kranti Meher, Madhuri Rotella, Subhramanyam Vangala, Satish Chandran, Nikhil Malhotra, Ratnakar Palakodeti, Sreedhara R Voleti, Uday Saxena

## Abstract

Drug repurposing can expedite the process of drug development by identifying known drugs which are effective against SARS-CoV-2. The RBD domain of SARS-CoV-2 Spike protein is a promising drug target due to its pivotal role in viral-host attachment. These specific structural domains can be targeted with small molecules or drug to disrupt the viral attachment to the host proteins. In this study, FDA approved Drugbank database were screened using a virtual screening approach and computational chemistry methods. Five drugs were short listed for further profiling based on docking score and binding energies. Further these selected drugs were tested for their in vitro biological activity. There was significant correlation between the prediction from computational studies and the actual RBD-ACE2 binding inhibition by the drugs. Then, we performed a series of studies that mimic some of the biological events seen in COVID-19 patients such as secretion of IL1β, presentation of a more thrombogenic endothelium by production of thrombomodulin and accumulation of inflammatory cells such as monocytes in the lungs. Of all the drugs, most promising drug was Ertugliflozin which is used for type-2 diabetes. This drug possesses several desired properties and may be a good candidate for immediate repurposing for treatment of COVID-19.

## 1. Introduction

World Health Organization has now declared COVID-19 a global pandemic actively spreading around the world. COVID-19 caused by the virus SARS-CoV-2, can cause symptoms such as fever, cough, pneumonia, nausea, and fatigue, and death due to severity of symptoms and associated co-morbidities such as hypertension, diabetes, etc. As of now, SARS-CoV-2 has reached almost all countries around the world, with more than 121 million confirmed cases and around 3 million deaths as of March, 2021 [1]. The mechanism of COVID-19 effect is believed to be that it binds its host’s cellular receptors through, homo viral surface trimeric spike glycoprotein (S-protein) with human host cell surface angiotensin-converting enzyme (hACE2) receptor and membrane fusion, necessary for viral entry into the host cell [2]. The entry of the virus into the host cell requires the spike or S-protein to be cleaved in two steps. The first step is the binding of the virus to the ACE2 protein and this is accomplished by the cellular proteases acting at the region between S1 and S2, namely, the transmembrane serine protease TMPRSS2. S1 and S2 subunits are responsible for viral-receptor binding and virus-host cell membrane fusion, respectively. The S1 subunit bears the receptor-binding domain (RBD) on its C-terminal domain. The RBD itself contains the receptor-binding motif (RBM), which actually comes into contact with the carboxypeptidase domain of the ACE2 molecule. In the spike protein, there is unique furin cleavage site which was not found in SARS-CoV. This site makes the virus more adaptable for cleavage by other cellular proteases as well, thus making it able to infect more cell types. This cleavage step is required for the initiation of membrane fusion by the S2 subunit, and this involves a second site of cleavage on this site, which is now considered essential to activate the S2 protein [3].

Mitigation of COVID-19 by drugs prophylactically and therapeutically has become the focus of pharma industry research. To this end, in early 2020, many anti-viral compounds were proposed [4], natural products were reported [5] and now, vaccines are being developed [6].

It is agreed universally that drug-repurposing is a safe and best approach towards treatment of COVID-19. Drug repurposing refers to the identification of novel applications for an approved or investigational/experimental drug outside the premise of its original indication. At present, this strategy would be a logical choice for developing a new drug for COVID-19 considering the substantial time-scales associated with new drug discovery and the trial-based validation of its safety and efficacy. The major advantage of repurposed drug is that it has been already evaluated for safety in human trials, which would reduce the significant amounts of time and money, a priority concern in SARS-CoV-2 drug development.

Application of in silico techniques in early stages of drug discovery either through conventional or repurposing approaches aid in minimizing the chances of the failures. The SARS-CoV-2 virus is closely related to the SARS-CoV and this allows utilization of the known protein structures to quickly build a model for drug discovery to search for possible medications for the SARS-CoV-2. The aim of the present study is to in silico identify clinically approved drugs treating various diseases of human importance, which would then be pharmacologically evaluated for targeting the RBD domain of the S-protein of COVID-19 as well as their anti-inflammatory and anti-thrombogenic properties.

The RBD domain of S-protein is an attractive and well characterised drug target in corona viruses owing to the pivotal role it plays in the entry of the virus into the host cell. Targeting this specific structural domain with repurposed drugs to disrupt the COVID-19 viral attachment to the host proteins. Much research has gone into finding drugs that can effectively prevent or reduce viral-ACE2 interactions. Subsequently, testing their binding efficacy in in vitro assays with RBD will elucidate the computational rationale for robust proposal of drug repurposing. Screening the FDA approved drugs and antivirals from the Drugbank database through molecular docking methodology would identify and prioritize repurposed drugs as potential leads in order to avoid any bias towards a particular drug. The drugs identified by virtual screening were then tested in a cell-free biochemical RBD-ACE2 binding study to validate findings from virtual screening. In addition, the drugs were further profiled in cytokine secretion and three dimensional (3D)-printed human-cell-based vascular lung model for potential anti-inflammatory and anti-thrombotic activities which could be useful in treatment of COVID-19.

## 2. Material and Methods

Cresset Flare software was used for molecular docking studies against the spike protein SARS-CoV-2 (http://www.cresset-group.com/flare/) [7].

### 2.1 Ligand preparation

The 2D structures of all the drugs were downloaded from Drugbank database and prepared using Flare software. Hydrogen atoms were added in the structure and atom force field parameterization was assigned. Further, energy minimized was done for all the drugs, nonpolar hydrogen atoms were merged, and rotatable bonds were defined. Later, ligand minimization has been carried out in Flare by Minimize tool by using Normal calculation methods. Ligands should also be prepared to assign proper bond orders and to generate the correct tautomer and/or ionization state.

### 2.2 RBD structure preparation

The RBD of spike glycoprotein SARS-CoV-2 is used for present study. This RBD binds to ACE2 receptor on the host cell with high affinity, which makes it a key target for the novel coronavirus therapy development. The 3D structure of RBD binds to ACE2 receptor (PDB ID: 6M0J) were downloaded from Protein Data Bank (PDB) (https://www.rcsb.org). The protein has two chain A and E, the A chain has ACE2 receptor and E chain has RBD domain. The RBD domain has been save in to PBD format for further studies. The target protein preparation was carried out in Flare software with default settings. Missing residues, hydrogen’s and 3D protonation were carried out on the target protein. Protein minimization has been carried out in Flare by Minimize tool by using Normal calculation methods.

### 2.3 Computational analysis of binding sites

Binding site was generated Accelrys Discovery Studio visualizer 3.5 (Copyright© 2005-12, Accelrys Software Inc.) to explore potential binding sites of the RBD protein using receptor cavities tools. Based on a grid search and “eraser” algorithm, the program defines where a binding site is. The binding sites were displayed as a set of points (point count) and the volume of each of cavity was calculated as the product of the number of site points and the cube of the grid spacing. Volume of each site were calculated and further saved and exported in to Flare for advance analysis.

### 2.4 Docking with Lead Finder

The full-atom model of RBD was prepared and docking of ligands to the prepared model of RBD in the active sites was performed using Flare (Lead Finder) software [8]. by default configuration setting. The energy grid box for ligand docking was set at the geometrical centre of the active site residues to span 10 Å in each direction. Two active sites were identified and in each sites centre residues were picked to define the active site grid box. Lead Finder assumes that the protein is rigid and analyses the possible conformations of the ligand by rotating functional groups along each freely rotatable bond. For each ligand pose Lead Finder determines values of the free energy of binding, the VS score, and the pose ranking score by using its three built-in scoring functions. Three different scoring functions are available from Lead Finder.

#### Rank Score

optimized to provide accurate prediction of 3D docked ligand poses.

#### ΔG

optimized to provide an accurate estimate of protein-ligand binding energy, on the assumption that the pose is correct.

#### VS

optimized to provide maximum efficiency in virtual screening experiments, with a maximum discrimination between active and inactive compounds in virtual screening experiments.

The database drugs were docked in the RBD domain of S-protein SARS-CoV-2 by using Cresset Flare Docking software with accurate and slow mode. The best poses were generated and visualized in pose viewer and 3D images stored in storyboard.

### 2.5 RBD-ACE2 biochemical assay

The above assay was performed using SARS-CoV-2 sVNT ready to use kit by Genscript, which is a competition ELISA, mirroring the viral neutralization process. In the first step, a mixture of horse radish peroxidase-RBD (HRP-RBD) and controls/drugs (drugs at the concentrations of 50µM, 100µM and 200µM) were incubated at 37°C for an hour to allow the binding of drugs to HRP-RBD. Following the incubation, these mixtures were added to a capture plate, which was a 96 well microplate coated with human ACE2 (hACE2) receptor to permit the binding of any unbound HRP-RBD and the ones bound to drugs to ACE2 receptor. After incubating the microplate at 37°C for an hour, plate was washed four times using wash buffer in order to remove any unbound circulating HRP-RBD drug complexes. Washing step was followed by addition of a colour substrate; tetramethyleneblue (TMB), turning the colour to blue. The reaction was allowed to run for 15 minutes followed by the quenching using stop solution turning the colour from blue to yellow. This final solution was read at 450 nm. The absorbance of the sample is inversely proportional to the inhibition of RBD’s binding to ACE2 by the drug. Data is presented as percent inhibition relative to untreated wells.

### 2.6 Interleukin-1β (IL-1β) secretion assay

Human IL-1β ELISA was performed to detect the presence of IL-1β which is a key mediator of inflammatory response. The ELISA was performed using Human IL-1β high sensitivity kit from Invitrogen. The sample of ELISA used in the present study consist of cell culture media (referred to as sample hereafter) collected after treating the cells with drugs at 100µM. Samples were added to the microplate precoated with human IL-1β antibody which captures the IL-1β present in the samples. A secondary anti-human IL-1β antibody conjugated to biotin was added to the plate. Following an overnight incubation, microplate was washed six times using wash buffer in order to remove any unbound biotin conjugated anti-human IL-1β antibody. Streptavidin-HRP was then added which binds to biotin conjugated antibody and the plate was incubated at room temperature on a shaker for an hour. After the incubation, plate was washed again following the same process as the previous wash step and an amplification reagent I was added to the wells. Following the incubation of 15 minutes and a wash, amplification reagent II was added. After incubation of half an hour in dark and a wash step later, a substrate solution was added turning the colour to blue. After 15-20 minutes, the reaction was terminated using a stop solution (turning the colour from blue to yellow). This final solution was read at 450nm. The OD of the sample is directly proportional to the amount of human IL-1ß present in it.

### 2.7 Human Thrombodulin/BDCA-3 Immunoassay

Human Thrombodulin ELISA was performed using a ready to use sandwich ELISA kit by R and D systems. Samples (media collected from our 3D vascular lung system) were added to a microplate precoated with monoclonal antibody specific for human thrombodulin. This was followed by an incubation of two hours (allowing any thrombodulin present in the sample to bind to the monoclonal antibody), the plate was washed with wash buffer four times. After washing an enzyme-linked monoclonal antibody specific for human thrombodulin was added to the plate. Another wash step was performed in order to remove any unbound antibody-enzyme reagent after completion of two hours of incubation. A colour substrate was then added turning the colour to blue. Reaction was quenched after 15-20 minutes using a stop solution which turned the colour from blue to yellow. Absorbance was read at 450 with wavelength correction set to 540 nm or 570 nm. Data is presented as amount of thrombomodulin in the media.

### 2.8 3D vascular lung model

#### Cell culture

For the 3D vascular lung model three types of cell were grown:

1. A549 cells were grown at 37o C in the growth medium DMEM (HIMEDIA #Cat No-AT007) supplemented with 10% (v/v) fetal bovine serum under the atmosphere containing 5% CO2. Cells were subcultured after reaching 80-90% confluence.
2. HUVEC cells were grown at 37o C in the Endothelial cell basal medium-2 (LONZA #Cat No-CC-3156 and CC-4176) under the atmosphere containing 5% CO2. Cells were subcultured after reaching 80-90% confluence.
3. HL60 cells were grown at 37o C in the growth medium RPMI (HIMEDIA #Cat No-AT028) supplemented with 10% (v/v) fetal bovine serum under the atmosphere containing 5% CO2. Cells were subcultured after reaching 80% -90% confluence.

### 2.9 3D bioprinting

In the 3D-vascular lung model four layers were bioprinted, first layer was collagen layer, (30µl of Rat tail collagen) was added to each well in 96 well plate and incubated for 1 hour in CO2 incubator at 37°C. After incubation, once the collagen is solidified A549 cells flask which has reached 80-90% confluence were trypsinized and cells were counted with the help of haemocytometer. Cell suspension was loaded into bioprinter syringe and A549 cells were printed in each well of 96 well plate and the cells were incubated for 48 hours.

After 48 hours of incubation, previous A549 media was removed carefully and again 30µl of Rat tail collagen coating solution was added to each well in 96 well plate and incubated for 1 hour in CO2 incubator at 37°C. After incubation, once the collagen is solidified HUVEC cells flask which has reached 80-90% confluence was trypsinized. Cell suspension was loaded into syringe and HUVEC cells were printed in each well of 96 well plate with help of 3D bioprinter and the cells were incubated for 48 hours.

After 48 hours of incubation the media was removed and the final selected drugs from virtual screening were added with LPS at 0.5 µg/ml and without LPS at the desired concentration. Later, cells were incubated overnight. Next day the drugs were removed and the cells were washed once with media. Endothelial cells were fixed with 4% paraformaldehyde for 3 minutes at 40°C. After fixing with paraformaldehyde, the cells were again washed with media.

MTT stained HL60 monocytic cells were added to each well in 96 well plate and incubated the plate for one hour and the wells were then washed twice with the media and pictures were taken of all the wells and the bound HL 60 cells were counted with the help of ImageJ, an open access software platform.

## 3. Results and Discussion

### 3.1 Structure of the S-protein

The S-protein is in a state of trimeric form with three RBDs and in metastable pre-fusion conformation that undergoes structural rearrangements to fuse the viral membrane with the host human cell membrane. The RBD undergo hinge-like conformational movements when attached with the ACE2 host receptor. The binding of virus particles to cell receptors on the surface of the host cell is the initiation of virus infection; therefore, receptor recognition is an important determinant of viral entry and a drug design target. The S-protein has two subunits S1 and S2, RBD situated in the S1 subunit binds to the cell receptor ACE2 in the region of aminopeptidase N. Full spike protein has 1,273 amino acids in length, and the amino terminus and most of the protein is predicted to be on the outside of the cell surface or the virus particles. A fragment that is located in the S1 subunit and spans amino acids 319–541 has the RBD domain. S1 protein has N-terminal domain (13-305 amino acid residues) and the RBD domain (amino acids 319-541), within the RBD domain there is one motif which makes complete contact with the receptor ACE2, was referred to as receptor-binding motif (RBM) (437-508 amino acid residues). The RBM region is tyrosine rich. Among the 14 residues of RBM that are in direct contact with ACE2, six are tyrosine, representing both the hydroxyl group and hydrophobic ring, as shown in **Figure-1**. S2 is responsible for membrane fusion and comprised of fusion peptide (FP), heptad repeat 1 (HR1), heptad repeat 2 (HR2), transmembrane domain (TM), and cytoplasmic domain fusion (CP), is responsible for mediating viral fusion and entry. For all COVIDs, S is further cleaved by host proteases at the so-called S2’ site located immediately upstream of the fusion peptide.

**Figure-1:**
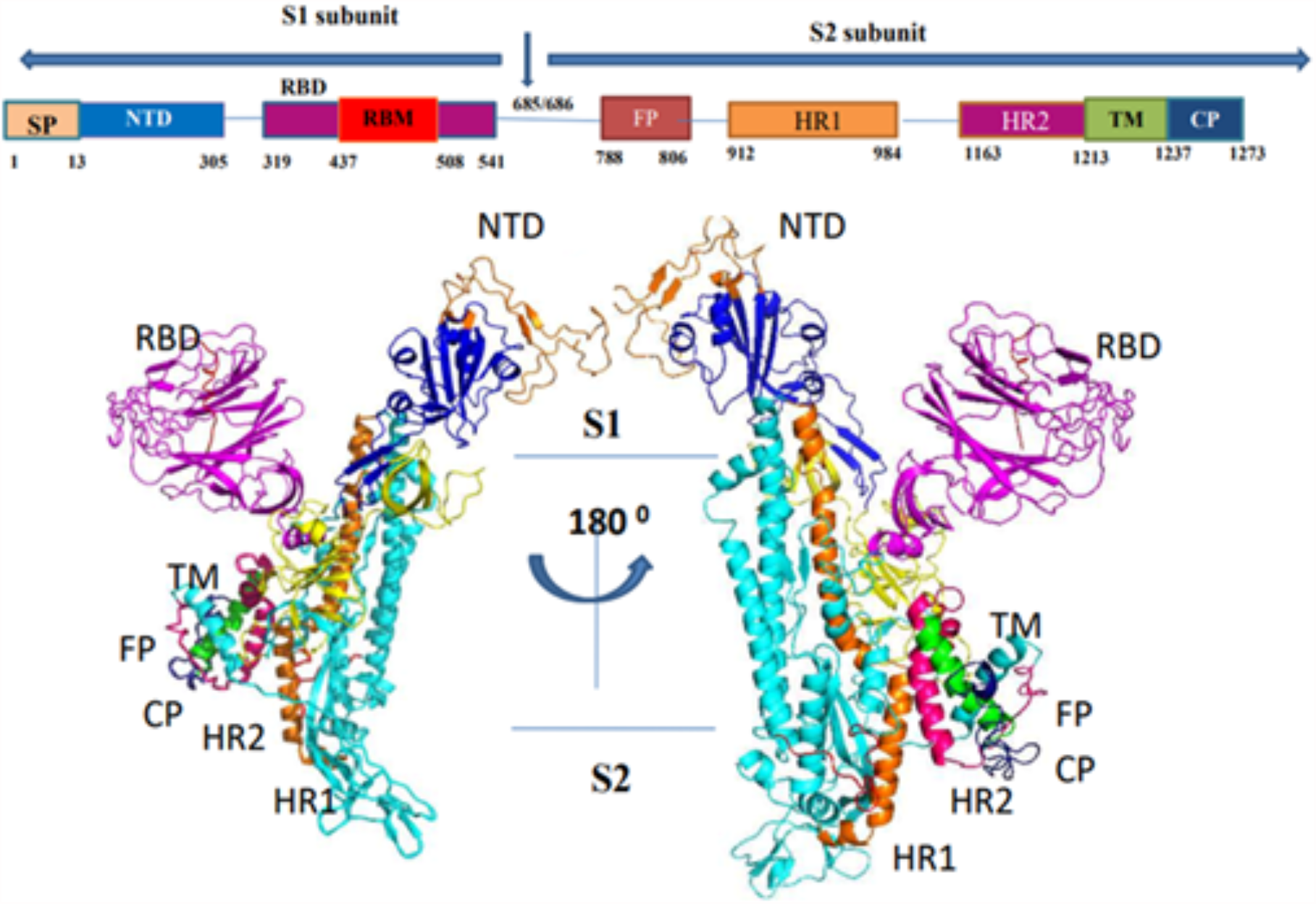
Monomer S-Protein-Structural Classification.

### 3.2 Identification of active sites

Our *in silico* strategy help us to design and screen the new drug targeting the S-protein that contains key structural domains RBD which plays a pivotal role in viral-host attachment. One such avenue of research is to find new repurposed drug that can attach to residues at the site of binding of the RBD to the ACE2. In the current study we had discover several potential binding sites for molecules that can occupy such druggable pockets so as to inhibit virus-ACE2 binding in vitro. The X-ray model of the RBD was used to identify 3 possibly druggable pockets where drugs might bind. The active site volume and binding surface area of three pocket is representation in **Table-1**. Site-1 (S1-1) has the volume of 143.7 & site-2 (S1-2) has the active site volume of 109.4, the last site-3 (S1-3) has active site volume of 87.46. Site-3 was too small to accommodate ligands, so they could not be potential pockets. Based on size of active site cavity site-1 & site-2 has been selected for further docking studies.

**Table-1:** Active sites of RBD

| Site | Binding surface area | Active site volume |
| --- | --- | --- |
| S1-1 | 673.6 | 143.7 |
| S1-2 | 668.3 | 109.4 |
| S1-3 | - | 87.46 |

### 3.3 Virtual Screening (VS) with Lead Finder (LF)

After the identification of active sites, the VS was applied which allows a large chemical library to be screened computationally against a specific drug target. All the database compounds were subjected to docking using LF and the predicted binding poses were analysed. In the present study, we used Structure-based Virtual Screening (SbVS) which uses molecular docking techniques to screen large virtual libraries of database where all the molecules were docked in the RBD domain of spike protein. The compounds are scored based on the predicted interactions with the target protein and those with the top scores (hits) are selected for further analysis. All the database drugs were docked in the spike protein SARS-CoV-2 (PDB ID: 6M0J E-chain) by using Flare Docking module of Cresset software. For all the database molecules less reasonable physicochemical parameters were discarded, leading to the selection of candidates with good drug-like properties. Later, a virtual library of approximately 3458 compounds extracted from Drugbank database were used for further screening using High-throughput virtual screening (HTVS) mode and top 100 molecules were finally docked in the “slow & accurate mode”. All compounds were ranked based on their docking score values and those with a score ≤ -8.0 and ligand having contact with more than one hydrogen bond in both the active sites were taken forward for final docking in “slow & accurate mode” of Cresset’s Flare Docking software.

### 3.4 Molecular Docking and Interaction Analysis

Docking analysis and visualization of virtual screening hit sled to the identification of several drugs that bind selectively to the RBD site-1 and site-2. In our previous studies, we have identified antiviral and kinase inhibitors as prospective targeted therapy for COVID-19 [9-10].

In this study rigorous *in silico* analysis through molecular docking analysis showed 5 drugs have strong predicted binding affinities toward RBD domain in the S1 protein. Total 5 drugs were identified based on rank score (RS), and binding energy (ΔG) and interaction with important amino acids in both the active sites of protein. These include unrelated and distinct set of approved drugs namely Levothyroxine, Iodipamide, Cromogilic acid, Iohexol and Ertugliflozin, which has shown most favourable binding affinity toward RBD S-protein.

The docking results showed that some drugs bound to RBD site-1 and some bound to RBD site-2. Our goal was to find those molecules which has great binding affinity towards both the active sites. Detail binding analysis of selected drugs towards active site of spike protein SARS-CoV-2 was studied in detail. Interaction analysis of drugs with spike protein SARS-CoV-2 (RBD) were carried out to identify the compound having highest binding affinity with target protein. Active site-1is encompasses of Arg454, Phe456, Arg457, Lys458, Glu465, Arg466, Asp467, Ile468, Ser469, Glu471, Thr473, Gln474 and Pro491 amino acid residues, while the site-2 encompasses the residues Leu335, Cys336, Pro337, Phe338, Gly339, Trp436, Phe342, Asn343, Val362, Ala363, Asp364, Val367, Leu368, Ser371, Ser373 and Phe374 amino acid residues, as shown in **Figure-2**. Active site-1 is more hydrophobic in nature as compare to active site-2.

**Figure-2:**
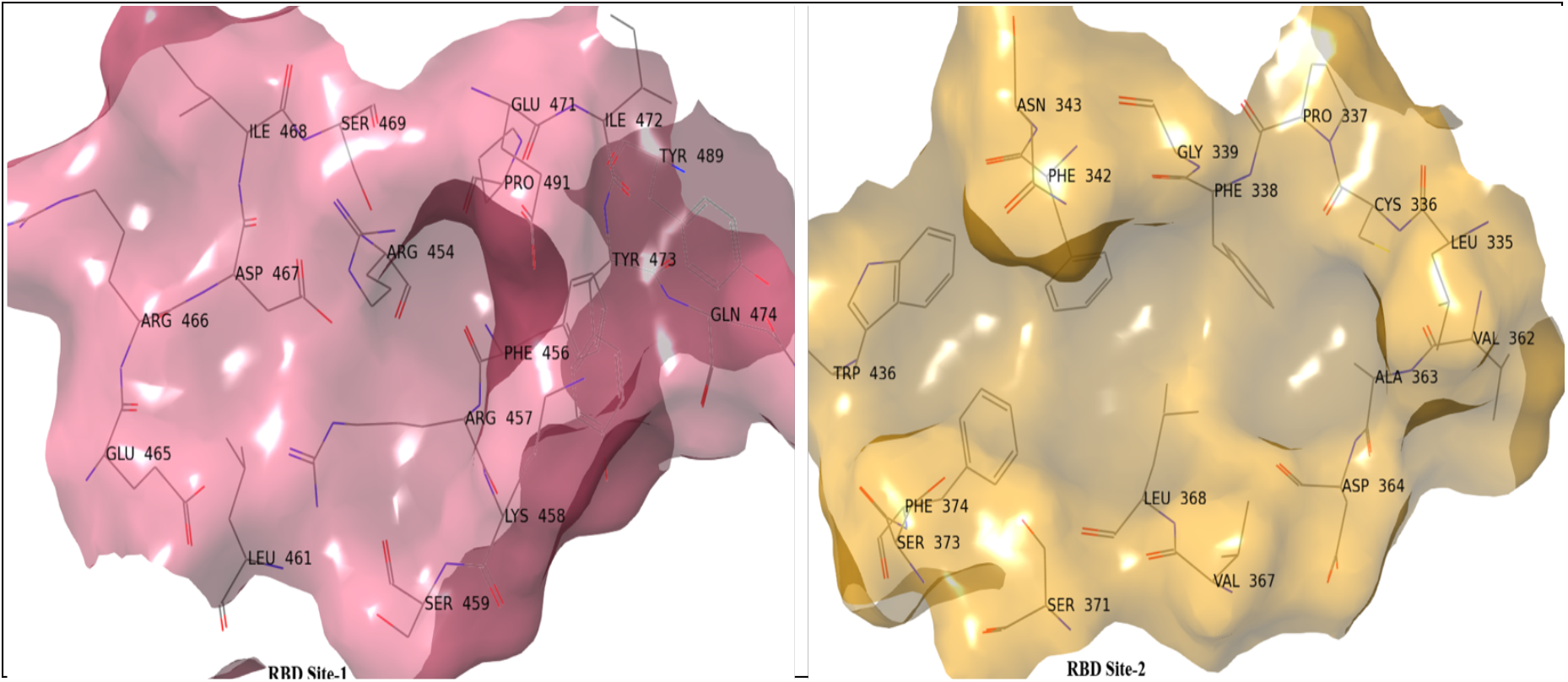
Active sites of RBD domain of S1-1 and S1-2 of Spike protein.

The LF rank score is an indicator of the binding affinity of protein-ligand complex. The LF rank for selected drugs in the pockets 1 & 2 is described in **Table-1 & 2** respectively. The binding orientation for each selected drugs having the least LF rank score, more negative LF rank score represent the better affinity of the drugs against target SARS-CoV-2 S-protein.

**Table-2:** Highest scoring molecules for SARS-CoV-2 Spike RBD in Site-1.

| S1-1 |  |  |  |
| --- | --- | --- | --- |
| FDA Drugs | $\Delta G$ | Rank score | Interactions |
| Cromoglicic acid | -7.51 | -11.62 | Arg454, Lys458, Asp467, Glu471 |
| Levothyroxine | -10.82 | -11.44 | Asp467, Ser469, Glu471, Gly482 |
| Iohexol | -9.13 | -10.22 | Arg457, Asn460, Glu465, Arg466, Asp467, Glu471 |
| Iodipamide | -9.51 | -10.73 | Arg457, Lys458, Arg466 |
| Ertugliflozin | -8.61 | -11.71 | Phe456, Lys458, Asp467, Ile468, Glu471 |

Among the docking studies performed on drugs, all the drugs had effective binding interactions with RBD domain of SARS-CoV-2 S-protein in both the active sites. The LF rank score values were in the range of -10.2 to -11.7 in the active site-1, while in active site-2 the LF rank score range from -8.8 to -11.7 for all the identified drugs. The ΔG provide an accurate estimate of protein-ligand binding energy, on the assumption that the pose is correct. For the selected drugs the ΔG is range from -7.5 to -10.8 in site-1, while in site-2 the ΔG is range from -7.6 to -10.8. The number of hydrogen bond and the number of amino acid residues of SARS-CoV-2 (site-1 & site-2) interacting with each drugs are given in **Table-2 &****Table**-3. As shown in **Figure-3**, the graph represents the correlation between rank score and binding energies of all the selected drugs in RBD site-1 and site-2. All the drugs have more or less similar range of score and binding energies.

**Table-3:** Highest scoring molecules for SARS-CoV-2 Spike RBD in Site-2.

| S1-2 |  |  |  |
| --- | --- | --- | --- |
| FDA Drugs | $\Delta G$ | Rank score | Interactions |
| Cromoglicic acid | -7.63 | -11.71 | Cys336, Phe338, Asn343, Ser373 |
| Levothyroxine | -10.72 | -11.44 | Cys336, Asp364, Ser371 |
| Iohexol | -9.72 | -8.84 | Phe342, Val367, Ser373, Arg509 |
| Iodipamide | -10.81 | -10.52 | Cys336, Phe342, Asn343, Trp436 |
| Ertugliflozin | -8.91 | -11.72 | Gly339, Asp364, Cys366, Arg509 |

**Figure-3:**
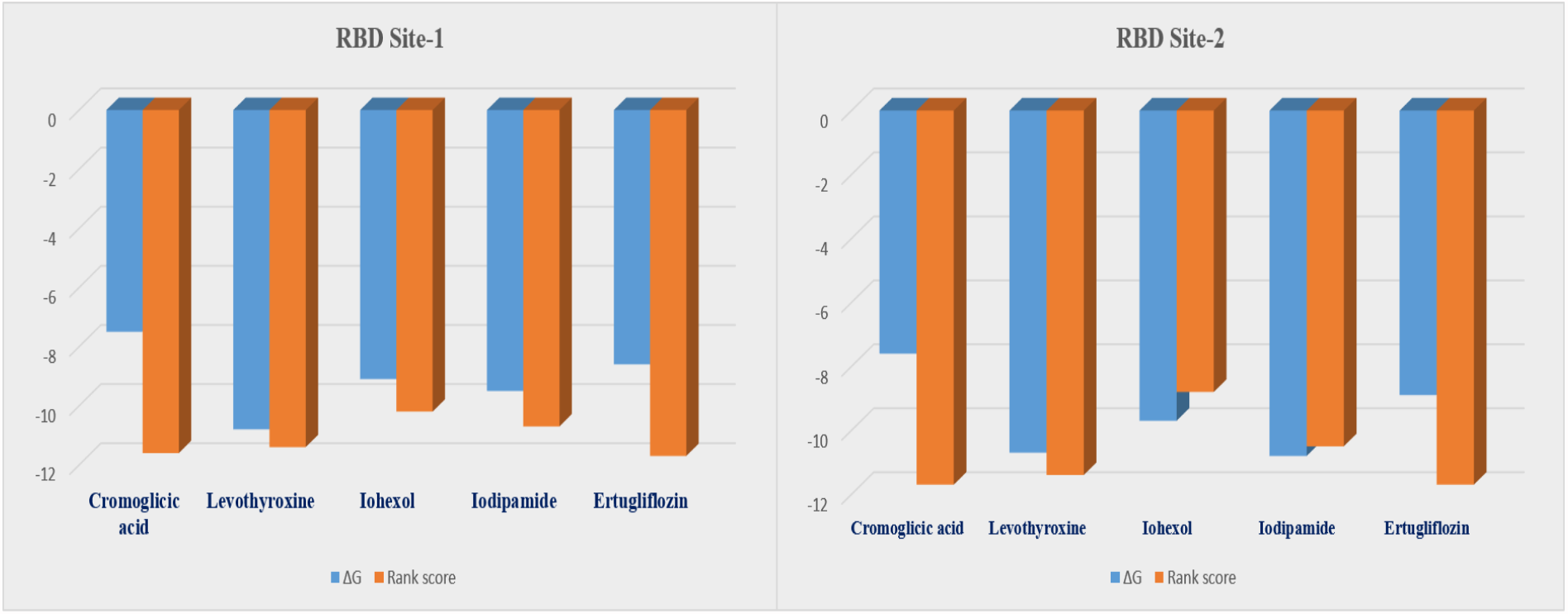
Rank score and binding energies of selected drug s in RBD’s site-1 & site-2.

From the detailed docking analysis, it is observed that the **Ertugliflozin and Iodipamide** drugs have good LF rank score and ΔG in both the active sites. It is found that, these six drugs have formed strong H-bond contacts with more than two to three amino acid residues along with various hydrophobic interactions in spike protein resulting in increased binding affinity with target protein. **Table-2 and Table-3** shows the rank score, binding energy, and list of interactions of the prioritized ligands with the site-1 and site-2 of the RBD.

Out of all the selected drugs, Ertugliflozin a Sodium-glucose cotransporter-2 inhibitor (SGLT2) inhibitor showed best free energy and rank score toward the binding to the RBD domain of spike protein of SARS-COV-2. SGLT-2 inhibitors, a new novel class of antidiabetics are used to manage glucose levels in Type 2 Diabetes, promotes excretion of glucose thus reduce plasma glucose levels. Our computational studies on the potential of various other SGLT-2 inhibitors binding to the RBD domains resulted either similar or lower LF rank scores. The LF rank score and ΔG for the other SGLT2 drugs in the RBD active site pockets S1-1 and S1-2 are described in **Table-4**. Based on these observations we believe that the effects of Ertugliflozin are unique for COVID-19.

**Table-4:**
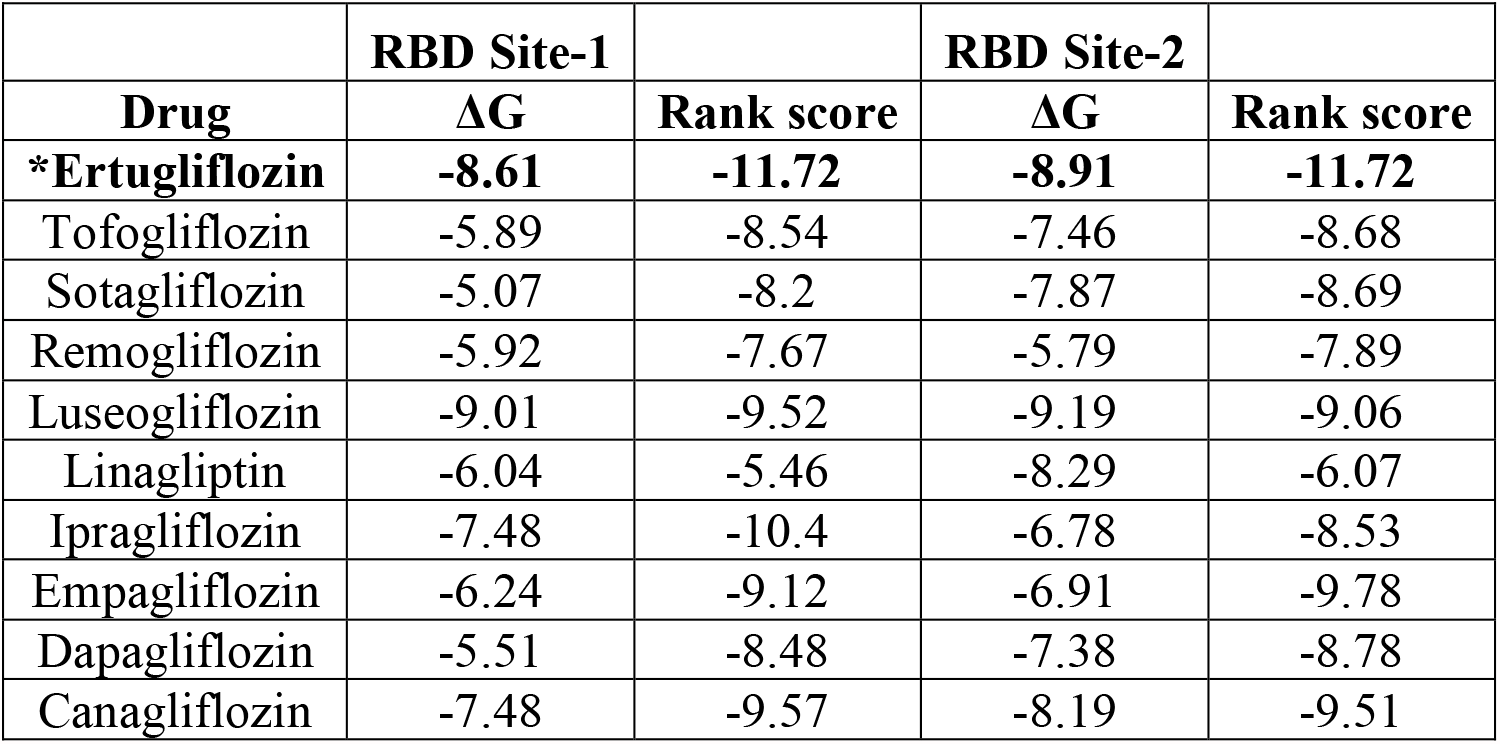
Docking score and binding energies of other SGTL2 inhibitors in SARS-CoV-2 Spike RBD in Site-1 and 2.

|  | RBD Site-1 |  | RBD Site-2 |  |
| --- | --- | --- | --- | --- |
| Drug | $\Delta G$ | Rank score | $\Delta G$ | Rank score |
| **\*Ertugliflozin** | **-8.61** | **-11.72** | **-8.91** | **-11.72** |
| Tofogliflozin | -5.89 | -8.54 | -7.46 | -8.68 |
| Sotagliflozin | -5.07 | -8.2 | -7.87 | -8.69 |
| Remogliflozin | -5.92 | -7.67 | -5.79 | -7.89 |
| Luseogliflozin | -9.01 | -9.52 | -9.19 | -9.06 |
| Linagliptin | -6.04 | -5.46 | -8.29 | -6.07 |
| Ipragliflozin | -7.48 | -10.4 | -6.78 | -8.53 |
| Empagliflozin | -6.24 | -9.12 | -6.91 | -9.78 |
| Dapagliflozin | -5.51 | -8.48 | -7.38 | -8.78 |
| Canagliflozin | -7.48 | -9.57 | -8.19 | -9.51 |

*Ertugliflozin* has demonstrated most favourable binding affinity toward the RBD protein in both the sites. Its belong to the class of potent and selective inhibitors of the Sodium-dependent Glucose cotransporters (SGLT), more specifically the type-2 which is responsible for about 90% of the glucose reabsorption from glomerulus. The docking orientation of Ertugliflozin in site-1 & site-2 is representation in **Figure-4**.

**Figure-4:**
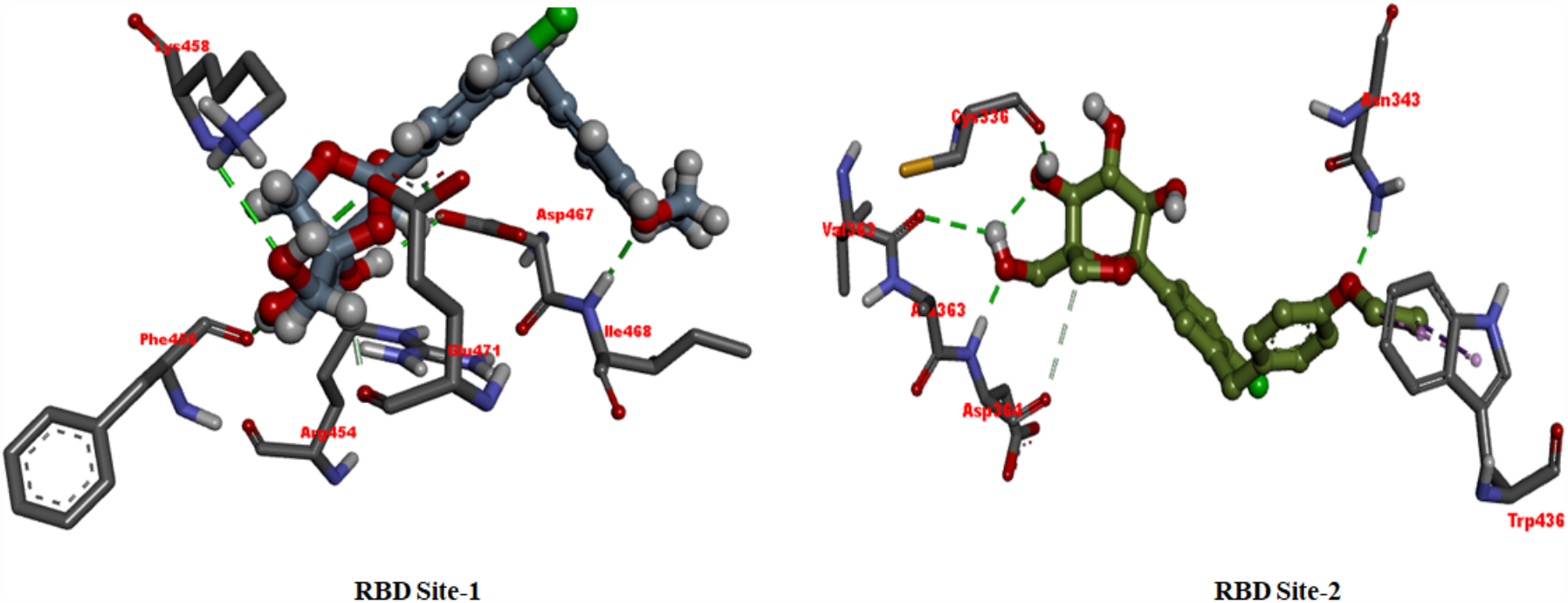
Binding orientation of Ertugliflozin within the RBD site-1 and site-2.

In site-1, Ertugliflozin showed conventional hydrogen bonds with Asp467, Ile468, Phe456 along with halogenic bond with Glu471 amino acid residue. The drug further stabilized by strong *cation-pi* interaction with Lys458. In site-2, Ertugliflozin is involved in hydrogen bonding interaction with Gly339, Asp364, Cys366, Arg509 amino acid residues. Additionally, it has *pi-pi* interactions with Phe374. The drug was found placed proximal to various hydrophobic amino acids such as Leu335, Trp436, Pro337, Phe338, Phe342, Val362, Ala363, Val367 and Leu368.

Another drug, Iodipamide, which is used as contrast agent for cholecystography and intravenous cholangiography is also showing good LF Rank score and ΔG in both the active sites. In site-1, it makes three hydrogen bonding interactions with Arg457, Lys458, and Arg466. Further, it also is involved in strong *cation-pi* interaction with Arg457. Interaction analysis in active site-2 suggests that it has four hydrogen bonding interactions with Cys336, Phe342, Asn343, Trp436 amino acid residues. Further it has making a close contact with various hydrophobic interactions with Leu368, Asn440, Phe347, Arg509, Val367, Ser371 and Ser373 amino acid residues, as shown in **Figure-5**.

**Figure-5:**
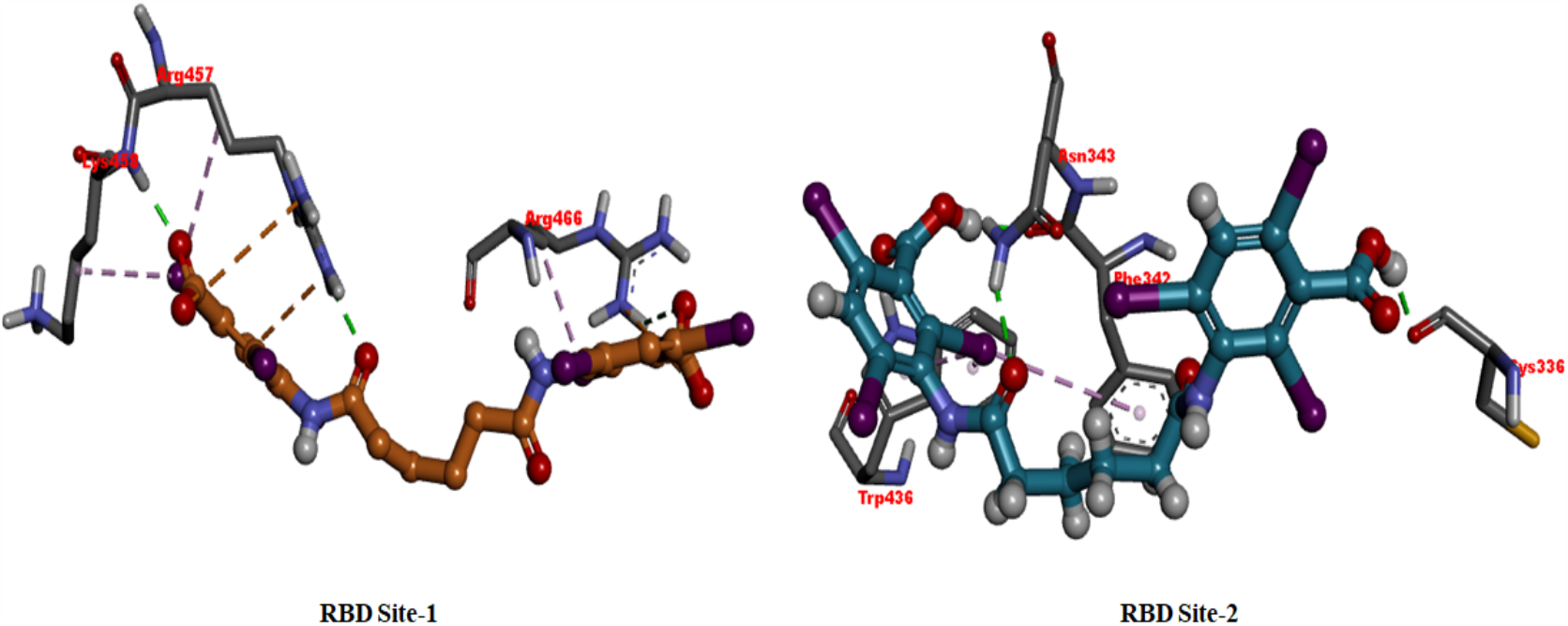
Binding orientation of Iodipamide within the active site of RBD site-1 and site-2.

### 3.5 Pharmacological evaluation of selected drugs in RBD-ACE2 binding assay

Top screened drugs from the *in silico* studies were evaluated biologically against RBD-ACE2 binding assay at various concentrations from 200µM to 50µM. In this ELISA assay, we have tested the drugs ability to block the binding of biotinylated RBD to ACE2. The data are shown as percent inhibition relative to binding observed in the absence of any drugs. As shown **Figure-6**, most drugs had some degree of inhibition thus corroborating the *in silico* predictions. Two drugs Iodipamide, Ertugliflozin showed excellent dose response inhibition that approached more than 90% inhibition of binding of RBD to ACE2 and correlated well with the *in silico* studies.

**Figure-6:**
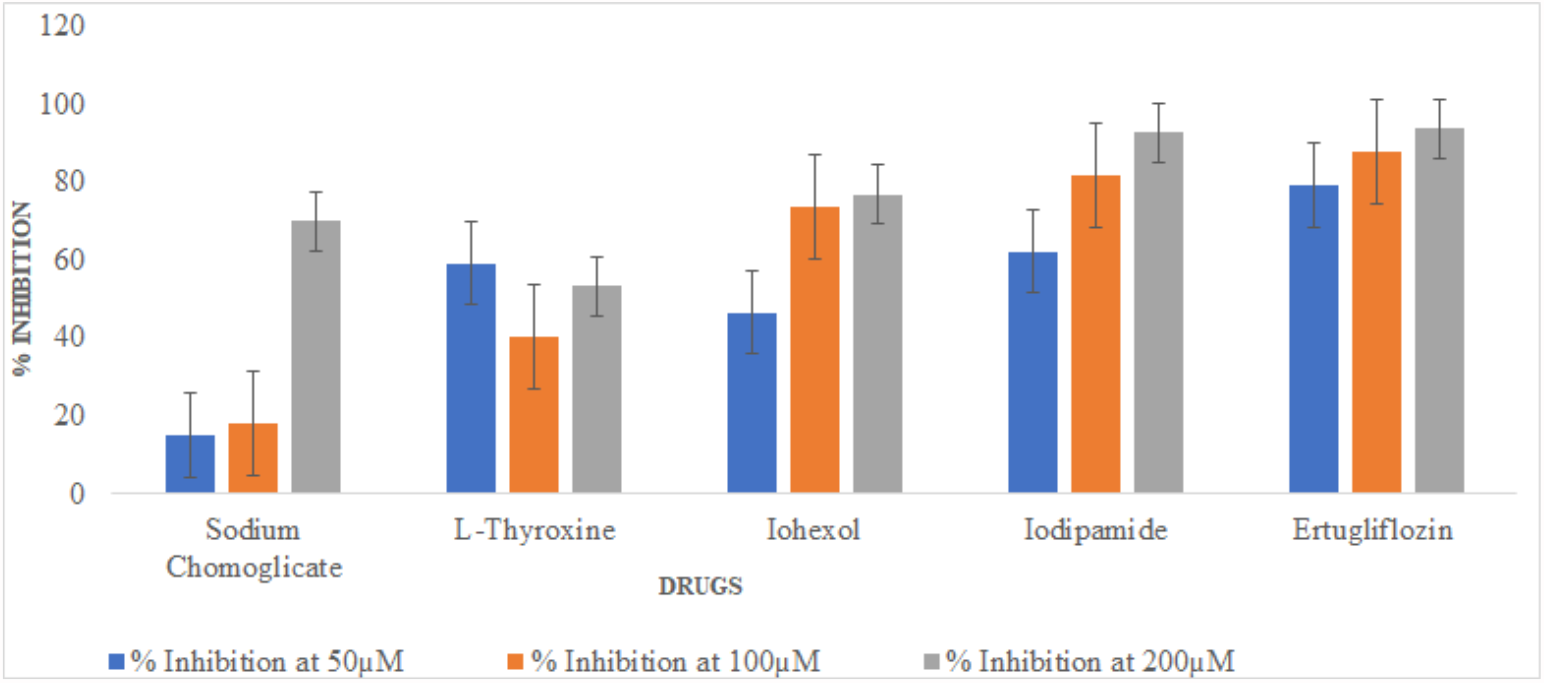
Percent Inhibition by different drugs at different concentrations of 50µM, 100µM and 200µM on RBD binding to ACE-2. *** P<0.001; ** P<0.01; *P<0.05. Student’s t-tests were performed to compare between percent inhibition shown by control and drugs.

### 3.6 Inhibition of IL-1β secretion

Further, we evaluated the effect of selected drugs on secretion of IL-1β a proinflammatory cytokines that are observed during cytokine storm observed in COVID-19 patients. To induce secretion of cytokines by the A549 lung cells, we used lipopolysaccharide/endotoxin (LPS) a known stimulus for inflammation and thermogenesis in many studies. As shown in **Figure-7**, there was a modest but significant increase in IL-1β secretion by LPS relative to untreated cells. When the drugs were added together with LPS some of them blunted the induced secretion. Almost all drugs showed moderate inhibition of LPS induced cytokine secretion. We became specially interested Ertugliflozin since it’s an approved drug for type-2 diabetes and has large body of safety data upon chronic use and has good potential for repurposing.

**Figure-7:**
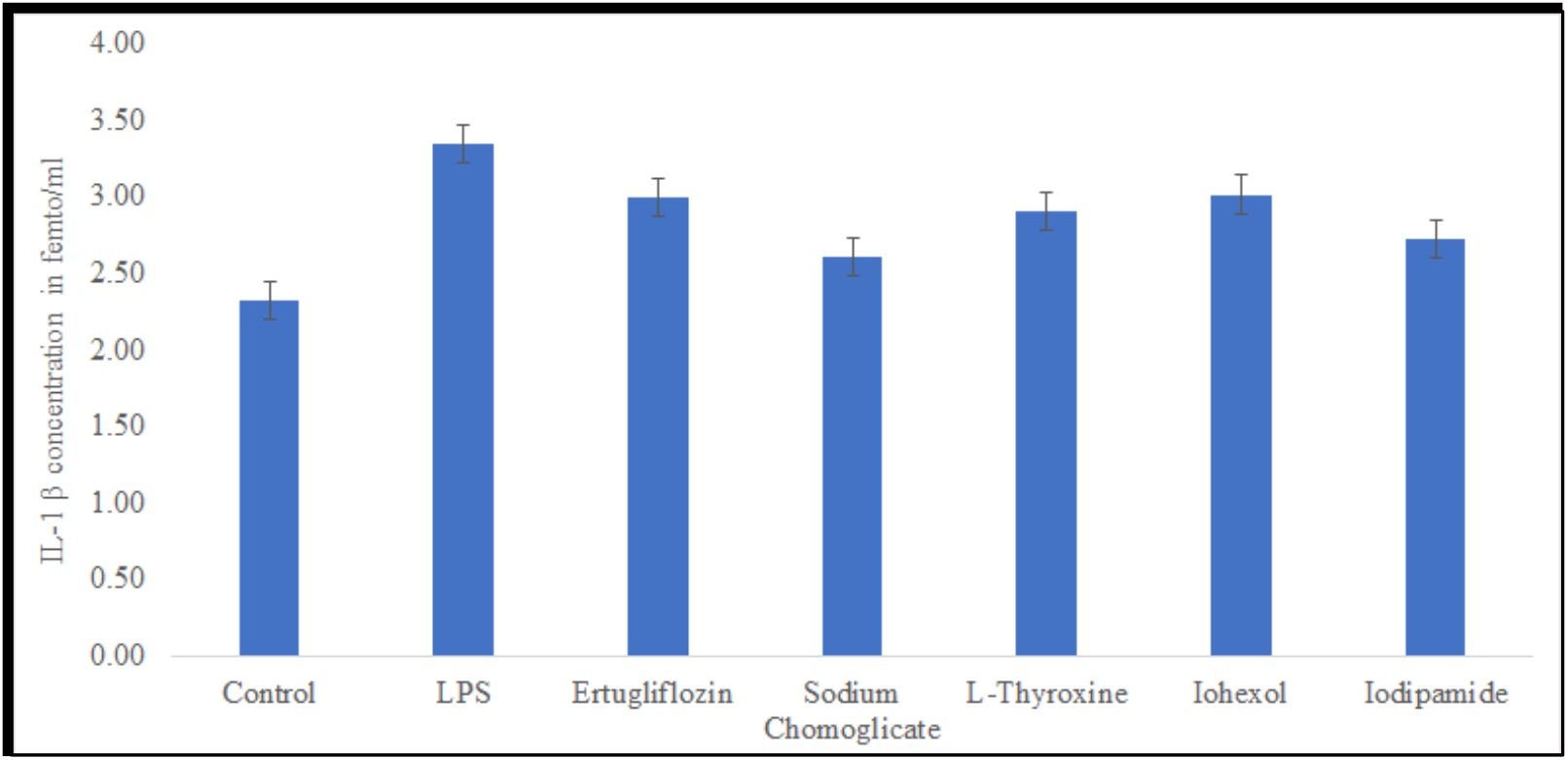
Effect on IL-1β secretion by different drugs (tested at 100µM). *** P<0.001; ** P<0.01; *P<0.05. Student’s t-tests were performed to compare IL-1β secretion between control and LPS treated cells; and between LPS treated cells (acting as a control) and different drug treated cells.

### 3.7 Effect of drugs on thrombodulin secretion, a procoagulant protein secreted by endothelial cells

We next looked at effect of the drugs on thrombomodulin a cell surface and secreted protein from endothelium which promotes blood clotting. We studied this protein since in more recent studies it has become obvious that uncontrolled blood clotting is seen in COVID-19 patients and could contribute to mortality. As reported in literature LPS induced the secretion of thrombomodulin in the 3D bioprinted vascular lung model. Almost all drugs had some degree of inhibition but continuing our interest in Ertugliflozin we were able to see that it blocks the secretion of thrombomodulin by almost 40%, a very desirable property to possess for a drug that can be repurposed for COVID-19 (**Figure-8**).

**Figure-8:**
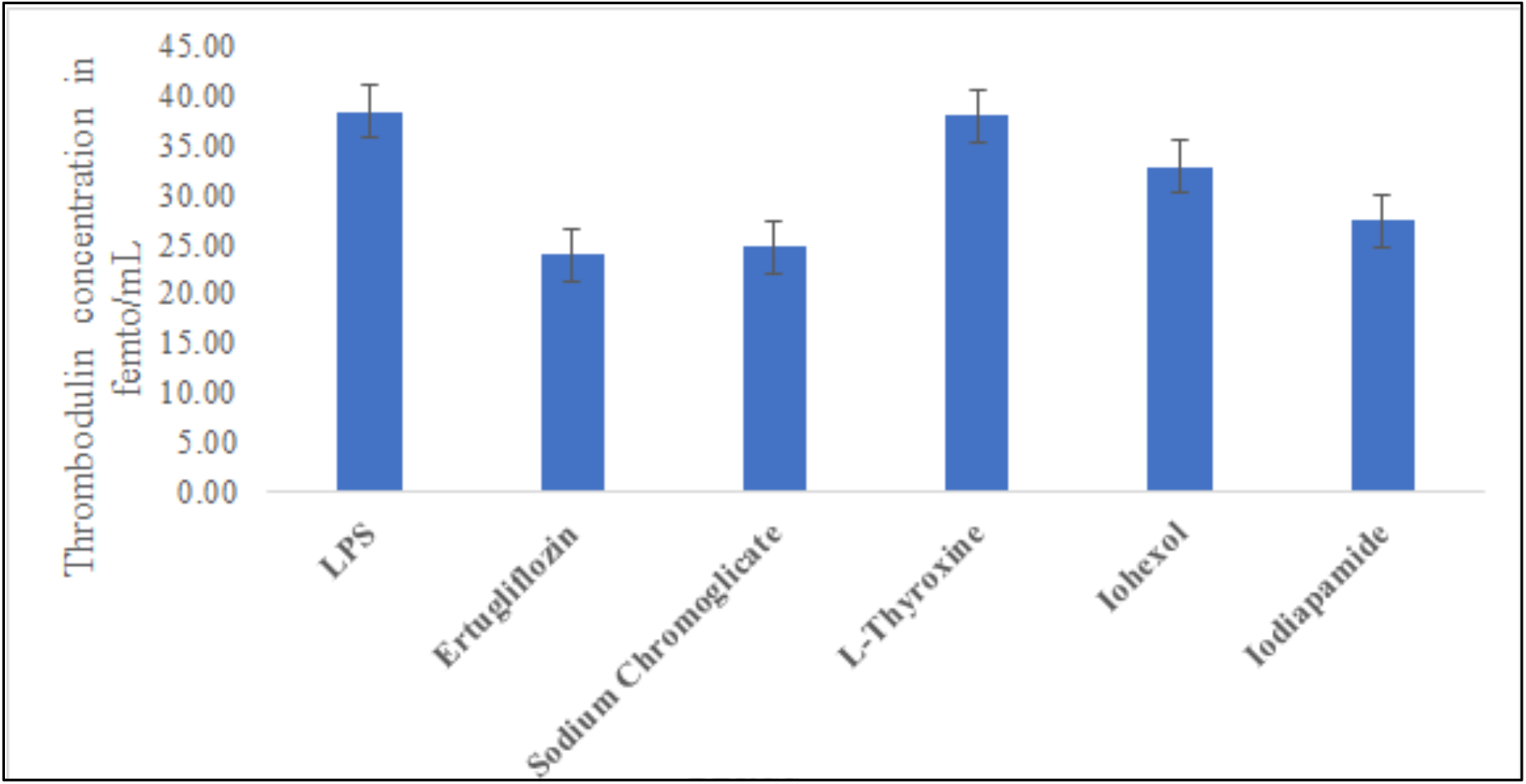
Effect on Thrombodulin secretion by different drugs (tested at 100µM concentration).

### 3.8 Effect of drugs on adhesion of monocytes

We finally examined the effect of drugs on adhesion of HL60 monocytes to endothelium in our vascular lung model. The recruitment of inflammatory cells such as monocyte is a common feature seen in COVID-19 as an early step of disease onset. This monocyte accumulation may greatly contribute to cytokine storm that is seen in this disease. As shown in **Figure-9**, some of the drugs were able to modulate monocyte adhesion stimulated by LPS. Again focusing on Ertugliflozin, it can be seen that it was able to reduce the impact of LPS (increase is 20 cell count per field) relative to the increase seen in control cells (increase about 70 cell counts per field). These data indicate Ertugliflozin has potential to modulate monocyte accumulation observed during COVID-19.

**Figure-9:**
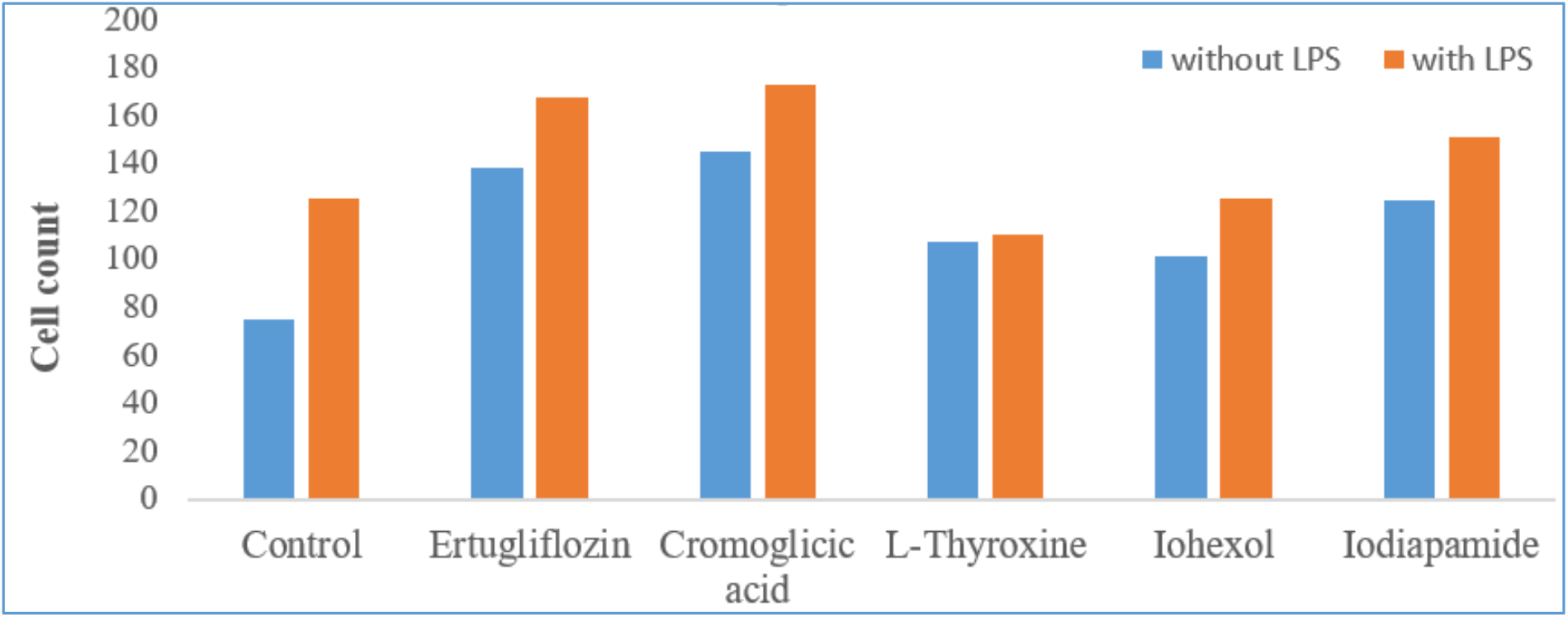
Effect on Monocyte adhesion by different drugs in 3D Vascular Lung Model. *** P<0.001; ** P<0.01; *P<0.05. Student’s t-tests were performed to compare between the control and drugs after treatment with LPS

There is an urgent need to identify a therapeutic for SARS-CoV-2 for blocking its spike protein binding to the host’s ACE-2, and quick way of finding such is through drug repurposing. RBD of S-protein plays a pivotal role in viral-host attachment and one of the potential target for antiviral treatment against SARS-CoV-2. These specific structural domains can be targeted with small molecules or drug to disrupt the viral attachment to the host proteins. In this study, FDA approved drugs were tested for their inhibitory properties towards the COVID-19 S protein (6M0J) using a virtual screening approach and computational chemistry methods. Rigorous *in silico* analysis through molecular docking analysis showed all 5 drugs have strong predicted binding affinities toward RBD domain in the S1 protein. All these also showed good dock score and interaction with important amino acids along with various hydrophobic interaction and were well fitted in both the active sites of protein. More importantly the selection of drugs by computational modelling was corroborated in a series of *in vitro* studies. The most important confirmation came from the assay that examined the direct binding of RBD to ACE2. There was very significant correlation between the prediction from computational studies and the actual binding assay. We then conducted a series of studies that mimic some of the biological events seen in COVID-19 such as secretion of IL-1β, presentation of a more thrombogenic endothelium by production of thrombomodulin and accumulation of inflammatory cells such as monocytes in the lungs. In silico and in vitro studies demonstrate favourably drug’s ability to modulate desirable properties, with Ertugliflozin standing out with promising potential for drug re-purposing.

## 4. Conclusion

Repurposing approved pharmaceutical drugs provide an alternative approach for rapid identification of potential drug leads. In the present study, we found Ertugliflozin, a drug used for type-2 diabetes, seems to possess all desirable properties demonstrated by in silico and in vitro pharmacological studies. This is an oral drug which works by blocking the activity of SGLT2 (Sodium-glucose co-transporter-2) and prevents glucose reabsorption in the kidney. This is used as a chronic treatment drug for diabetes, which means that its safety has been thoroughly tested prior to approval. It is also readily available and could be repurposed immediately. To avoid any glucose lowering in COVID-19 patients by this drug, a nasal formulation can be developed which targets the lungs. This will also improve the potency and efficacy against the infection while requiring lower doses. While the development of vaccines will prophylactically prevent the infection we do not have targeted drugs once the disease already sets in. Based on our work here we propose that Ertugliflozin is a strong candidate for immediate repurposing for treatment of COVID-19 treatment. Since the in silico screening did not identify other SGLT2 inhibitors in initial screening, we believe that the effects of Ertugliflozin are unique to it and unlikely to be a class effect, but this needs to be examined in separate studies. In conclusion, we propose that the disruption of the SARS CoV-2-RBD/ACE2 binding interface by Ertugliflozin and its associated anti-inflammatory and anti-thrombotic properties could pave the way for it to be useful as a new COVID-19 treatment.

## Acknowledgments

Dr. Uday Saxena would like to dedicate this manuscript in the loving memory of his father

Dr. E.R Saxena, his mentor Dr. K. Anji Reddy, his PhD advisor Dr. Sailen Mookerjea, while

Dr. Sreedhara Voleti in loving memory of his PhD advisor late Prof. A.K. Chandra.

## References

1. Hall Jr, D. C., & Ji, H. F. (2020). A search for medications to treat COVID-19 via in silico molecular docking models of the SARS-CoV-2 spike glycoprotein and 3CL protease. Travel medicine and infectious disease, 35, 101646.

2. Lan, J., Ge, J., Yu, J., Shan, S., Zhou, H., Fan, S., … & Wang, X. (2020). Structure of the SARS-CoV-2 spike receptor-binding domain bound to the ACE2 receptor. Nature, 581(7807), 215–220.

3. Huang, Y., Yang, C., Xu, X. F., Xu, W., & Liu, S. W. (2020). Structural and functional properties of SARS-CoV-2 spike protein: potential antivirus drug development for COVID-19. Acta Pharmacologica Sinica, 41(9), 1141–1149.

4. WHO Solidarity Trial Consortium. “Repurposed antiviral drugs for COVID-19-interim WHO Solidarity trial results.” New England Journal of Medicine 384.6 (2021): 497–511.

5. Vougogiannopoulou, K., Corona, A., Tramontano, E., Alexis, M. N., & Skaltsounis, A. L. (2021). Natural and Nature-Derived Products Targeting Human Coronaviruses. Molecules, 26(2), 448.

6. Kyriakidis, N. C., López-Cortés, A., González, E. V., Grimaldos, A. B., & Prado, E. O. (2021). SARS-CoV-2 vaccines strategies: a comprehensive review of phase 3 candidates. npj Vaccines, 6(1), 1–17.

7. Flare, version, Cresset®, Litlington, Cambridgeshire, UK; http://www.cresset-group.com/flare/

8. Lead Finder, version, BioMolTech®, Toronto, Ontario, Canada; http://www.cresset-group.com/lead-finder/

9. Saxena, S., Meher, K., Rotella, M., Vangala, S., Chandran, S., Malhotra, N., Voleti, S., & Saxena, U. (2021). In silico and in vitro Demonstration of Homoharrintonine Antagonism of RBD-ACE2 Binding and its Anti-inflammatory and anti-thrombogenic Properties in a 3D human vascular lung model. bioRxiv. doi: https://doi.org/10.1101/2021.05.13.443955

10. Saxena, S., Meher, K., Rotella, M., Vangala, S., Chandran, S., Malhotra, N., Voleti, S., & Saxena, U. (2021). Tyrosine Kinase Inhibitor Family of Drugs as Prospective Targeted Therapy for COVID-19 Based on In Silico And 3D-Human Vascular Lung Model Studies. bioRxiv. doi: https://doi.org/10.1101/2021.05.02.442384

